# Probabilistic Multi-site MR Image Harmonization via Feature Preserving Conditional Generative Adversarial Networks

**DOI:** 10.64898/2025.12.29.696930

**Authors:** Saeed Moazami, Sepideh Rezvani, Agnimitra Dasgupta, Assad A. Oberai

## Abstract

Brain magnetic resonance imaging (MRI) is pivotal in diagnosing and monitoring neurological disorders. However, despite their extensive applications, MR images have certain shortcomings. In particular, factors other than the anatomy of brain tissues influence the intensity distribution of voxels in MR images. These factors include hardware, software, magnetic field strength, and acquisition protocol. This inconsistency poses challenges in multi-site neuroimaging studies, where images are obtained from various devices with minimal control over acquisition parameters. Image harmonization algorithms aim to eliminate non-biological characteristics in MR images through various approaches, including converting images from multiple sites into a format resembling that of a designated target site. Among image harmonization methods, those relying on deep learning algorithms have gained significant attention recently. Nevertheless, certain aspects of deep learning-based image harmonization remain unexplored, notably the integration of probabilistic deep generative models to transform the distribution of MR images to a desired distribution. Inspired by this, we introduced a feature preserving conditional generative adversarial network (FP-cGAN) that converts images from multiple origins into the format of a target site while preserving anatomical features by imposing a novel regularizing constraint. We conduct our experiments on MR images from the SRPBS dataset, which comprises unpaired images in addition to paired (traveling subjects) images from multiple sites. We utilize the unpaired data for training our models and the paired data for evaluation. Furthermore, we compare our results with cycleGAN and histogram matching, two widely used image harmonization methods. Our experiments reveal that our approach surpasses the other techniques.

**Article Highlights:**

- We introduce an FP-cGAN model to harmonize multi-site MRI scans into a target site.
- A novel anatomical feature invariant constraint maintains tissue integrity.
- The models are trained in an unpaired setting without requiring traveling subjects.
- We evaluate harmonization performance using paired traveling subjects.

## 1 Introduction

Brain magnetic resonance imaging (MRI) is an instrumental, non-invasive medical imaging technique that offers detailed information about the internal structures and pathologies of the brain. It serves as the primary imaging technique for diagnosing and monitoring numerous neurodegenerative disorders, such as Multiple Sclerosis (MS), Alzheimer’s Dementia (AD), etc. MRI protocols are also highly flexible in providing a range of MRI sub-modalities, such as structural MRI, functional MRI (fMRI), and magnetic resonance angiography (MRA). Within each sub-modality, MRI sequences can be further tailored to offer a variety of MR image types or contrasts, like T1, T2, and fluid-attenuated inversion recovery (FLAIR), which are structural MR images. Each of these images serves a unique purpose in revealing distinct responses to various tissues, organs, and abnormalities [1, 2].

Despite the numerous advantages and widespread applications, MR imaging also suffers from inherent drawbacks. One significant limitation is the lack of a concrete ground truth, which can critically impact quantitative analyses that rely on MRI-based measurements. Theoretically, numerical values in MR images aim to represent the average of measured physical quantities within each voxel. However, these values are heavily influenced by the measurement process. Factors such as magnetic field strength, hardware setup, signal processing steps, pulse sequence design, and scanner drift (a gradual signal degradation and voxel intensity change over time [3]) can all impact the reported measurements. Additionally, noise induced by various factors, including hardware imperfections, thermal noise, and magnetic field inhomogeneity, distorts the measurements. These confounding factors can be categorized into known (e.g., site ID) and unknown sources of variation, such as scanner drift. As a result, MRI measurements yield a noisy representation of relative differences between tissues rather than exact values for specific physical tissue properties. This lack of consistency is often called the “batch effect” in the literature, implying the impact of the scanning device or batch. We also refer to this phenomenon as MR image variability or heterogeneity. MR image variability is considered a significant concern in MRI-based neuroimaging studies as it can lead to inaccurate results when automated computational tools use MR images as inputs [4]. Consequently, it can substantially compromise generalizability and reproducibility in downstream tasks. This issue is notably detrimental in multi-site neuroimaging studies, as the MR images are collected from multiple resources (inter-dataset variability). Multi-site studies are critical as they enable collaboration between centers to collect more data to improve the statistical power of the study and expand the target population. However, MR images in multi-site studies tend to demonstrate significant variability because it is challenging to maintain the same image acquisition parameters across all participant sites. We note that MR images within a dataset, even with consistent parameters, also exhibit inherent inhomogeneity (intra-dataset variability) [5].

Several studies have focused on quantifying the effect of MR image variability in downstream tasks [6–13], and a wide range of techniques have been introduced to account for this variability and mitigate the batch effect problem. These techniques are collectively referred to as image harmonization and come in two major categories. Techniques in the first category focus on harmonizing the results of tasks performed on MR images rather than the images themselves [14–16]. This includes harmonizing metrics like tissue volumes and thicknesses computed from MR images.

Techniques in the second category include MRI image harmonization methods that adjust voxel intensities across scanners, with early approaches correcting image histograms [17, 18] and more advanced ones using image-to-image transformations learned through deep learning. Paired-data methods train supervised models, such as U-Nets or diffusion models, on images of the same subjects scanned across sites, but these are limited by the scarcity and cost of traveling-subject datasets [19]. Other approaches manipulate latent space representations, disentangling anatomical content from scanner-dependent style using autoencoders or VAEs, though they face challenges in clearly defining and preserving biological information [20]. Generative adversarial network (GAN)–based methods, especially cycleGANs, harmonize images by enforcing style adaptation while preserving anatomical features through cycle consistency, though balancing adversarial and consistency losses remains difficult [21]. The proposed method builds on these ideas by employing a conditional GAN that combines adversarial loss, ensuring alignment with target distributions, and a structural preservation loss to maintain anatomical fidelity, producing harmonized MR images that reflect both style adaptation and biological accuracy.

The rest of the paper is organized as follows. In Section 2, we describe current MR image harmonization methods and also introduce the proposed approach. Thereafter, in Section 3, we describe this method in detail. Next, we demonstrate the results of our experiments in Section 4 and examine the effectiveness of our model. We provide conclusions derived from our study and suggest a number of possible future works in Section 5.

## 2 Image-to-image MRI Image Harmonization

In image-to-image MRI harmonization, a transformation is learned to map source images into their harmonized counterparts. Such mappings typically use deep learning models, which effectively capture complex relationships in high-dimensional image data. Figures 1 (a to d) schematically illustrate three representative image-to-image harmonization approaches alongside our proposed method. In these diagrams, blue and green points represent source and target MR images; circles and stars denote training and test data, respectively; arrows indicate learned transformations; and filled versus hollow points distinguish actual samples from model-generated images.

**Fig. 1:**
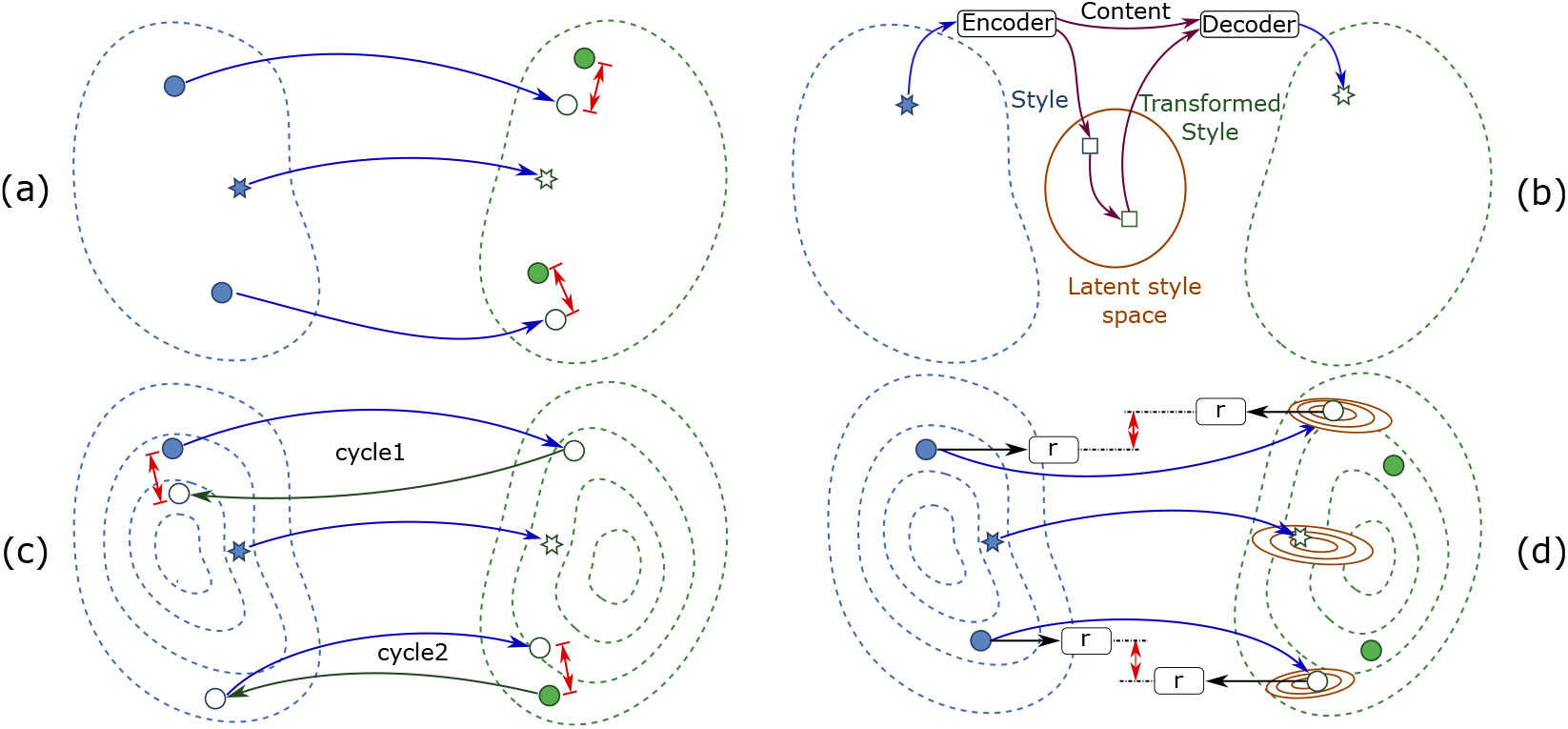
a) The schematic diagram for supervised learning-based image harmonization using paired data. The model learns the transformation from the source to target images by minimizing a difference loss. b) Image harmonization through latent space manipulation. An encoder transforms the input into a latent space containing content (geometry) and style (contrast). The style component is then transformed to match the target images. A decoder generates the harmonized image from the transformed style and unaltered content. c) Unpaired image harmonization using cycleGAN-based models. Two pairs of generators and discriminator models are trained using adversarial loss. The core idea is that when images are transformed from one site to another and back to the initial site, the output of this cycle should be the input. d) Our method uses adversarial loss to transform images to the target distribution. Also, a function (***r***) is used to calculate a measure of anatomical features to represent the geometry and keep it unchanged during transformation. Additionally, the model learns the distribution of likely outputs per given input (brown ellipses).

The first group of methods assumes access to paired images from traveling subjects or overlapping cohorts. These are scans of the same individuals across different sites, providing a ground-truth correspondence between source and target domains. These methods often rely on training a deep learning model using direct inference supervised learning methods to learn a transformation from the source to the target site, as shown by the blue curved arrows in Figure 1-a. The model generates transformed images (hollow green circles), and the loss (red double arrows) measures their deviation from the target images. Minimizing this loss yields transformations that closely match the ground truth. Along these lines, in [22], the authors utilize a U-Net [23] with a mean absolute error (MAE) loss function for 2.5D cross-protocol harmonization, where “2.5D” refers to combining axial, coronal, and sagittal slices to produce a 3D output. Paired-data harmonization is not limited to direct supervised inference but also extends to conditional generative models. For instance, the study presented in [24] utilizes denoising diffusion probabilistic models (DDPM) [25, 26] to learn conditional transformations between sites using paired data. While supervised and conditional approaches generally outperform unpaired (unsupervised/unconditional) methods [27], they depend on the availability of traveling-subject data. In practice, however, recruiting sufficient subjects with the necessary characteristics to scan across all participating sites is often infeasible.

Another group of methods, as illustrated in Figure 1-b, performs MRI harmonization via latent-space manipulation. Typically, encoder-decoder architectures such as autoencoders (AE) or variational autoencoders (VAE) map input images to latent vectors disentangled into anatomical features (content) and site-specific style. Harmonization is achieved by modifying the style while preserving content, and the resulting latent vector is decoded back into image space. Variants differ in architecture and algorithms for encoding, decoding, and disentanglement, and can operate on paired or unpaired data [19, 20, 28]. For example, the work presented in [28] employs a conditional VAE with intra-site paired T1 and T2 images to disentangle anatomy from style, enabling unsupervised inter-site harmonization of T1 images. This approach leverages multi-contrast intra-site data, which is easier to acquire than identical contrasts across multiple sites; however, such paired data is not always available. A general limitation of disentanglement methods is the ambiguity in defining content versus style and the difficulty of quantifying potential alterations to biological information during transformation.

The next group performs style or distribution matching directly in the native space of MR images using generative adversarial networks (GANs) [29–33] or combinations of GANs and VAEs [34]. During harmonizing, the transformed image must both belong to the target distribution and preserve the anatomical features of the input image. To this end, several methods employ cycleGANs [35] or other cyclic consistency variants [29–33]. CycleGANs typically consist of two generator-discriminator pairs trained concurrently: one maps images from source to target (blue arrows in Figure 1-c) and the other performs the reverse mapping (green arrows). The core principle of cyclic consistency is that an image converted from source to target and back should reconstruct the original input (cycle 1), and similarly for target to source and back (cycle 2). This consistency is usually imposed as an *L*_1_ loss (red double arrows in Figure 1-c).

In practice, cycle consistency has been observed to help with preserving the anatomical features in the transformed images, though we are not aware of a theoretical analysis of this feature. Utilizing cycle consistency also involves certain considerations. As noted in [21, 33], the relative weighting of the cycle consistency term against the adversarial loss requires careful tuning. Too small a weight biases the generated images toward the reference domain, while too large a weight preserves the source structure but limits adaptation to the reference style. We have also included cycleGAN models in our comparisons presented in this work.

Our image harmonization method takes a distinct approach compared to the techniques outlined above. Specifically, we employ a conditional GAN with an adversarial loss that enforces the transformed source images to align with the target distribution, while an anatomical feature–preserving loss (detailed in Section 3) constrains the transformation to maintain structural fidelity. As shown in Figure 1-d, each source image is mapped to target-domain counterparts that both conform to the target distribution and preserve the anatomical features of the original input with high fidelity. These components are further elaborated in the following sections.

## 3 Theory and calculation

In this section, we describe the image harmonization problem formulation and propose the FP-cGAN model to solve it. We also discuss the dataset used for training and testing of the models, the evaluation strategies and metrics, the pre/post processing steps, and the methods used for comparison against our model.

### 3.1 MRI image harmonization problem formulation

Let *M* be the number of sites and 𝒮 = *{*1, *…*, *M}* the set of all sites. The source and target sites are denoted by *s* and *t*, respectively, where *s, t ∈ 𝒮*. Also, let ***X***_*i*_ denote a random vector that represents unpaired axial head MR images from the *i*-th site. 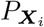 denotes the distribution of ***X***_*i*_ and ***x***_*i*_ represents a realization (sample) of ***X***_*i*_ sampled from 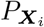. Images of all sites have the same number of pixels *d*_*X*_, and therefore 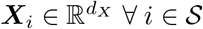.

We assume that for any *s, t ∈ 𝒮* with *s ≠ t*, there exists a function 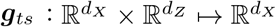, that takes as input realizations of 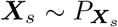 and random vector ***Z*** *~ P*_***Z***_, and maps them to a realization of 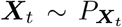. More formally, the target distribution 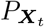 the push forward of the joint distribution 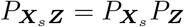 due to the map ***g***_*ts*_, i.e., 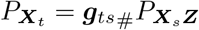. The random vector 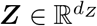, with its realization ***z***, is referred to as the latent variable and drawn from a simple distribution like a multivariate standard normal distribution, i.e., *P*_***Z***_ = 𝒩 (**0, *I***), where the covariance matrix 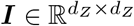 is the identity matrix.

We also assume that there exists a function 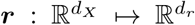 such that for any image sample 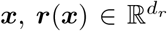 is a vector that retains information about the anatomical features of ***x***. We use ***r*** to enforce anatomical consistency during image harmonization. More specifically, we expect the push-forward ***g***_*ts*_ to be invariant with respect to anatomical features, i.e., for any realization of the input image ***x***_*s*_ and latent vector ***z***, we want ***r***(***x***_*s*_) = ***r***(***g***_*ts*_(***x***_*s*_, ***z***)).

We note that there is neither a universally accepted notion of anatomical features or geometry, in the context of MR images, nor a mathematical definition for the function ***r***. So, in this work, we propose ***r*** to be a function that yields the gradient of an image normalized by its Euclidean norm, i.e.,

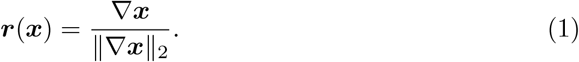

We remark that though we represent an image ***x*** in vector form, any image ***x*** will be a 2D array matrix in practice. So, we compute the gradient 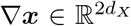 using finite difference approximations along the horizontal (*h*) and vertical (*v*) directions at each pixel location [*j, k*], i.e.,

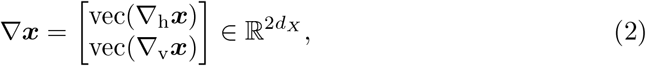

Where

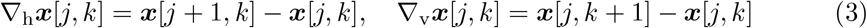

and vec(*·*) denotes the vectorization operator. Therefore, *d*_*r*_ = 2*d*_*X*_ as per the definitions above. Also,

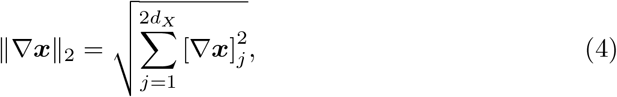

where [*∇****x***]_*j*_ denotes the j^th^ component of *∇****x***.

The motivation for using the gradient of an MR image as the representation of its anatomical features is the following. The gradient of the image captures the difference between adjacent pixels, which are unaffected by the absolute values of pixels. Also, normalizing the gradients forces the model to focus on preserving the direction in which changes in intensities occur instead of the magnitude of the changes. These properties are critical in the image harmonization setup, where the goal is to generate images with consistent structural content (e.g., tissue edges) across sites, while allowing intensity distributions to vary.

We aim to train ***g***_*ts*_ in a generative adversarial framework, using a regularization term to bound the changes of ***r*** due to the transformation induced by ***g***_*ts*_ and ensure that anatomical features are preserved. We will discuss this next.

In Figure 2, we demonstrate the proposed FP-cGAN architecture, which consists of two deep neural networks: a generator ***g***_*ts*_ and a critic *d*_*ts*_. The generator is designed to approximate the mapping function ***g***_*ts*_ described in Section 3.1. Given instances of input images and latent variables, the generator produces multiple transformed images 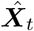 following the distribution 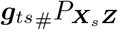, representing the pushforward of the joint distribution of the inputs 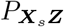 through ***g***_*ts*_.

**Fig. 2:**
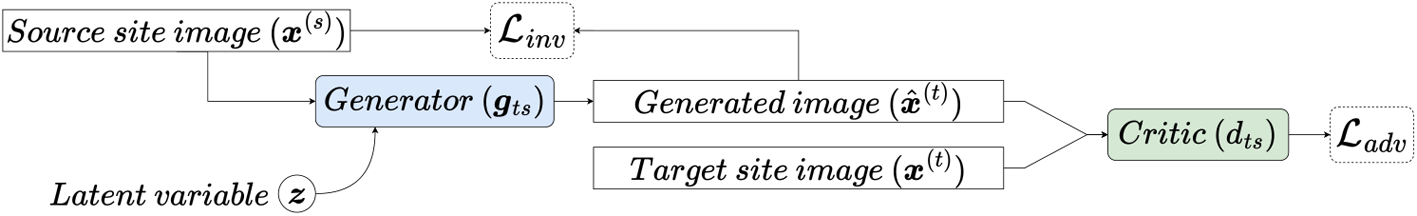
The feature preserving conditional generative adversarial network (FP-cGAN) architecture used in the proposed model. The generator ***g***_*ts*_ receives instances of the source images ***x***_*s*_ and generates images 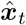 for random latent vectors ***z***. The input and output of the generator are constrained using an invariance loss term ℒ_*inv*_ to preserve the anatomical features during transformation. The critic *d*_*ts*_ distinguishes between the generated images 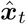 and ***x***_*t*_, i.e., images that are sampled from the target distribution.

The critic 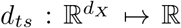 receives an image and is trained to distinguish between the inputs that are from the true target dataset, i.e., 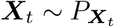, and those that are generated by the generator network, i.e., 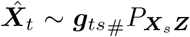. Through the training process, the critic yields larger values for images from the dataset (true) and smaller values for the images generated by the generator (fake). This is done by application of the Wasserstein GAN adversarial loss function [36] as the expectation of the difference between the values of the critic for true and generated images

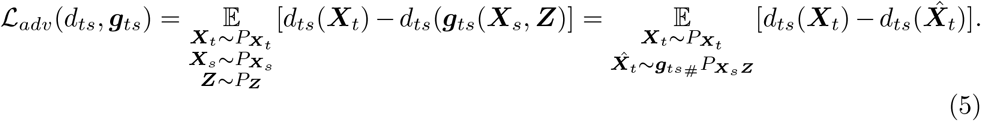

Then, the min-max optimization problem

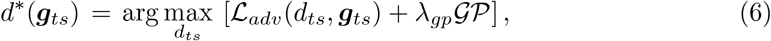

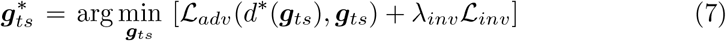

is solved concurrently to find the optimal generator 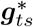 and critic 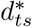. As we will discuss subsequently, ℒ_*inv*_ in Equation 7 is the anatomical features invariance loss term with coefficient *λ*_*inv*_ that makes use of the function ***r*** discussed in Section 3.1 to preserve the anatomical features during the transformation. Also, the term 𝒢 𝒫 in Equation 6 is the gradient penalty [3], with *λ*_*gp*_ coefficient, defined as

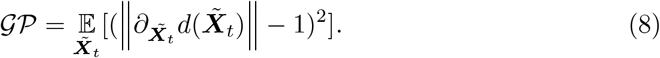

Intuitively, based on Equation 6, the fully trained optimal critic 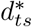 is the function that produces the greatest value for the loss objective defined in Equation 5 for all possible functions of ***g***_*ts*_ and *d*_*ts*_. The gradient penalty term in Equation 8 is used to enforce the 1-Lipshitz constraint to the critic by penalizing the gradients with absolute values larger than one (see [37] for details). The gradients are calculated with respect to 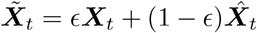, which is an average of actual and generated images, where the averaging is weighted using a random number *ϵ ~ 𝒰* (0, 1). After being trained, the optimal critic 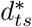 provides an estimate of the Wasserstein-1 distance between the two distributions of true and generated images. Training 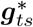 through Equation 7 is equivalent to finding a generator that minimizes this distance. Therefore, passing a given image ***x***_*s*_ with multiple instances of random latent vectors ***z*** through the fully trained generator 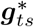 is equivalent to generating multiple samples from the true target distribution.

Additionally, ℒ_*inv*_ must be employed to enforce consistency during the transformation. This is necessary because an infinite set of transformations can couple the input and output distributions, and each of those may yield non-unique anatomical features or tissue geometries. Among them, we wish to choose a transformation that will preserve the features ***r***(***x***_*s*_). To this end, we define a distance function *ℓ* as the mean squared error (*MSE*) of ***r*** between two arbitrary images, say ***x***^(1)^ and ***x***^(2)^, as *ℓ*(***x***^(1)^, ***x***^(2)^) = *MSE*(***r***(***x***^(1)^), ***r***(***x***^(2)^)). Then, we utilize this function to define an anatomical invariance loss as the distance between the input and output of the generator as

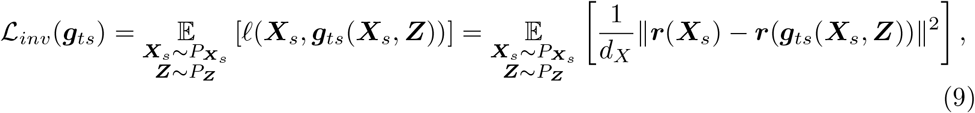

which is minimized jointly with the objective defined in Equation 7.

We illustrate the effect of this distance function using the blue curve in Figure 3. In this experiment, we compute the function *ℓ* used in the definition of ℒ_*inv*_ between a fixed image (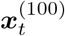: axial target image at slice *z* = 100) from a traveling subject scanned at a target site *t*, and a set of images from varying slices of the same subject scanned at a different source site and aligned with the target image, (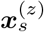:axial images from site *s* at slices *z∈ {* 1, …, 200*}*). As can be seen, this value is significantly lower when two images are from the same slice (at *z* = 100), i.e., they share the same anatomy, and remains high for slices with mismatched features. Accordingly, minimizing this distance as a loss acts as a regularization term during the training, guiding the generator to produce outputs that preserve the anatomical features of the input image during the transformation. We also illustrate the calculated Euclidean distance (*l*_2_) between the same set of images as a commonly used measure in the red curve in Figure 3. As can be seen, the introduced function *ℓ* shows significantly higher sensitivity to anatomy mismatch when compared to *l*_2_. We note that the curves are normalized to provide better visual comparison.

**Fig. 3:**
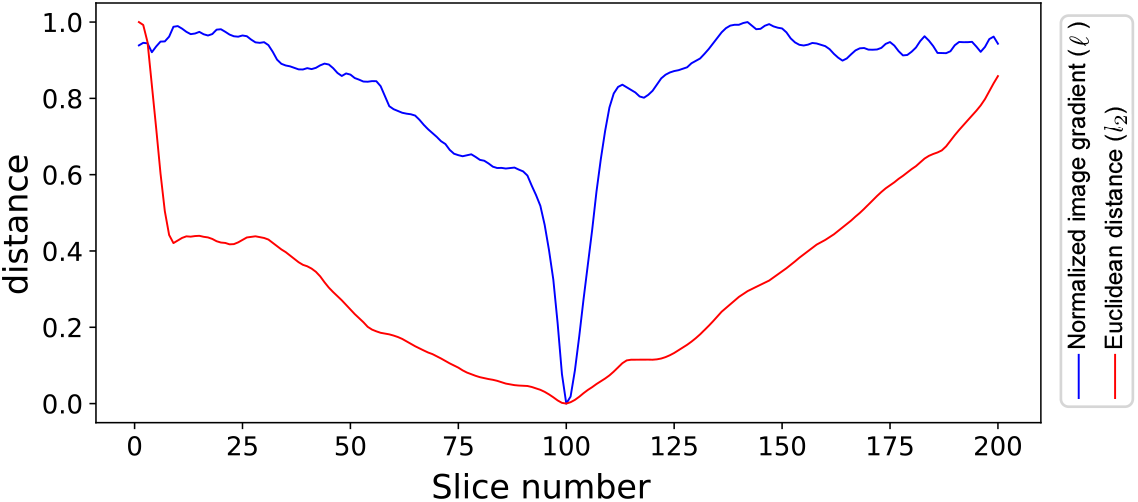
The blue curve shows the normalized image gradient function (*ℓ*) that captures the anatomical differences between a fixed slice image from a target site 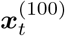 and a set of images from varying slices of the same subject, but scanned at a different site 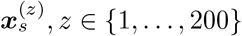, *z* ∈ {1, …, 200}. When the two images share the same anatomy (at *z* = 100), the calculated distance is significantly lower, and the distance remains high in mismatched slices. The red curve shows the calculated Euclidean distance (*l*_2_) between the same set of image slices. The function *ℓ* demonstrates higher sensitivity to mismatches of anatomical features.

As we discussed, the proposed FP-cGAN is capable of providing multiple outputs for any given input. Importantly, the generator model ***g***_*ts*_ requires only a single forward pass to generate an output, which makes this process computationally efficient. During the inference process, an input image ***x***_*s*_, which is an axial MR image of the head from the source site *s*, is passed to the fully trained generator along with *n* random instances of ***z***, yielding an ensemble of samples 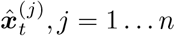. In this work, we calculate the final output 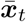 as the pixel-wise mean of the generated images as

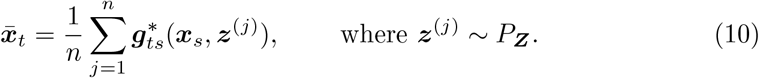

That is, the intensity of any pixel of a single computed transformed image 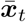 is the average of the values of the same pixel in *n* sampled output images.

### 3.2 Comparing methods

In addition to the proposed FP-cGAN model, we include Histogram matching, also referred to as Hist-match for brevity in this work, and cycleGAN as two commonly used methods from the image harmonization literature in our comparisons. Histogram matching is a method traditionally considered in the context of image harmonization [17, 18]. In this method, no training is required, and the intensity of the voxels is altered using a mapping so that the cumulative histogram of the output image matches the reference or a validation image. Comparing the results of histogram matching against our method provides insights into the quantitative performance and qualitative characteristics of the transformation that takes place during image harmonization (see Section 4 for examples). We note that handling MR artifacts, especially correcting the bias field, is critical before performing histogram matching. This is because histogram matching can further unveil the intensity inhomogeneity caused by field bias in MR images. We use the N4 bias field correction [38] tool [39] (an improved version over the N3 algorithm [40]) prior to histogram matching. We also include the cycleGAN model in our comparison as a common model in MR image harmonization literature when using unpaired data (see Section 1). For this purpose, we used the enhanced implementation [41] of the work presented in [35] using the original hyperparameters and the same training dataset as the FP-cGAN model.

### 3.3 Dataset

We use the SRPBS dataset [42] for training, validation, and evaluation of the proposed model^1^. This dataset comprises four sub-datasets from which we use SRPBS-Open for training and SRPBS-Traveling Subjects for validation and testing. The training dataset contains unpaired T1 MR images collected at 11 sites from 1410 patients at 3T. The Traveling dataset contains nine subjects who have undergone MR imaging at 12 participating scanners. In this study, we include a subset of sites with unique acquisition parameters that overlap both the Open and Traveling datasets. We use one subject for validation and eight subjects for testing and evaluation from the traveling dataset. Table 1 provides a summary of the characteristics of the sites used in this study.

**Table 1:**
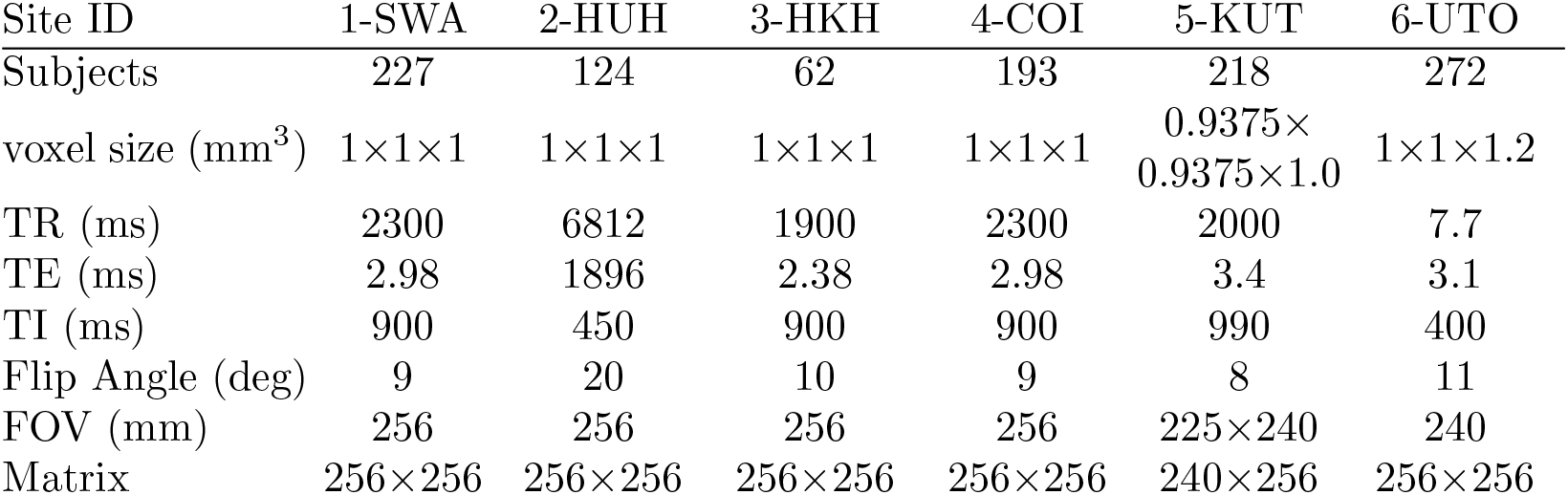
A summary of characteristics, including imaging protocols of different sites from the SRPBS dataset [42] used in this study. The following are the abbreviations used in the table: TR: repetition time, TE: echo time, TI: inversion time, and FOV: field of view.

As we discussed, the proposed model requires a target site to which it harmonizes the images from other sites. We note that there is no strict quantitative definition for the target site selection procedure. In practice, the target site can be selected following a set of rules depending on the study. For example, the target site should have a sufficiently large population size, high image quality, low intra-dataset heterogeneity, and a low number of artifacts, particularly hyper-intense outlier voxels and bias field. We observed that site 1-SAW has these characteristics. Therefore, we trained five models to transform images from source sites (2-UTO, 3-HKH, 4-COI, 5-KUT, and 6-HUH) to the target site (1-SWA).

### 3.4 Pre-processing

We aligned all the training images to the MNI-152 (0.5 mm) template volume [43] for spatial normalization using the FSL [44] FLIRT module [45] and scaled the axial slices to 240 × 200 pixels. For the traveling subjects, we first aligned the target images from site 1-SWA to the template volume. Then, we aligned the images from the source sites to the corresponding aligned target image for each subject. We also performed voxel intensity normalization, as different MRI devices produce images in various numerical data ranges. Accordingly, we performed a histogram mode matching as the first step. That is, we calculated the average of the values at which the image histograms attain their peak values. We then scaled the intensity of individual images so that they all have the same peak value. In the next step, we eliminated the hyper-intense outlier voxels of each image. For this purpose, we calculated a clipping value for all the images and set the values that are higher than the clipping value to the clipping value. Finally, we performed min-max normalization to ensure that all voxel intensities lie within the range of zero and one. We note that the pre-processing procedure should be followed separately for each site.

### 3.5 Evaluation and metrics

We evaluate the performance of the proposed FP-cGAN image harmonization method by applying it to the images of traveling subjects from the source sites (2-UTO, 3-HKH, 4-COI, 5-KUT, and 6-HUH) and comparing the outputs against the corresponding target images of the target site (1-SWA). We also include the comparison between the target images and the input images (before harmonization) and the outputs of Hist-match and cycleGAN. In our experiments, we incorporate the brain region of images by applying brain extraction [46] to the MR images, where we refer to the target brain image as ***Y*** and the test image (input or the outputs of harmonization methods) as ***B***. The availability of the traveling subjects enables us to utilize image-to-image similarity metrics, namely, the structural similarity index measure (SSIM) and peak signal-to-noise ratio (PSNR). The maximum possible value for SSIM is one, which indicates a perfect match between the target ***Y*** and the test image or prediction ***B***. We note that an SSIM of one is not practically achievable due to realignment mismatch (see Section 3.4) and the inherent variability of MR images. The latter implies that even if we scan a specific subject twice during one visit, the two images will not be exactly the same. Therefore, even a hypothetically perfect transformation can demonstrate slight differences from the target image.

## 4 Results and Discussion

In this section, we present the image harmonization results of the proposed FP-cGAN model and compare its performance against the Hist-match and cycleGAN methods. We also discuss neuroimaging observations and interpretations when appropriate.

In Figure 4, we showcase the harmonization results for an axial slice from the traveling subjects scanned at different sites. As indicated, each of the first five rows contains the images from a source site (2-HUH, 3-HKH, 4-COI, 5-KUT, and 6-UTO). The first column shows the input image. The second column is the absolute difference between the input and the target image. The target image is from the same subject scanned at site 1-SWA and is shown in the last row at the lowermost of the third, fifth, and seventh columns as a reference. The rest of the images in these columns (third, fifth, and seventh) are the image harmonization results of the Hist-match, cycleGAN, and the cGAN model for each corresponding source site. The columns to their right side (fourth, sixth, and eighth columns) are the output errors of each method, shown next to the corresponding outputs.

**Fig. 4:**
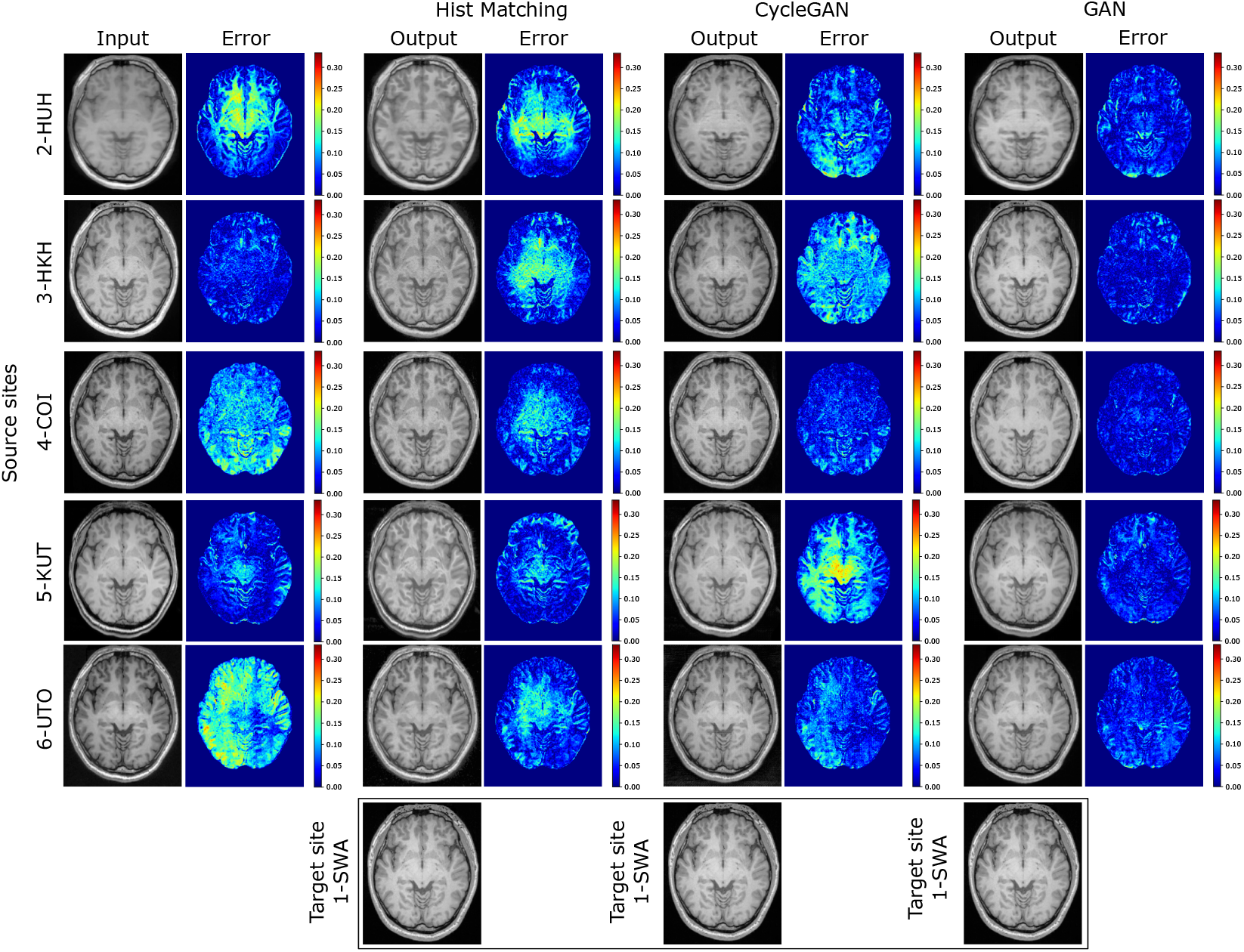
Image harmonization results of a typical axial image for different sites using histogram matching, cycleGAN, and FP-cGAN (ours). Each of the first five rows belongs to a source site. In each row, from left to right, the input image from the indicated source site and the error (absolute difference from the target image) are shown. The third, fifth, and seventh columns demonstrate the output of histogram matching, cycleGAN, and our FP-cGAN approach. The corresponding error is shown immediately to the right. In the last row, the target image is shown below the output columns of the three methods as a reference.

As can be observed from the sixth and eighth columns of Figure 4, the outputs of Hist-matching and cycleGAN demonstrate areas of relatively large errors. The inaccuracies occur both globally throughout the brain region, as seen in cycleGAN outputs for 3-HKH and 5-KUT (the second and fourth rows), and locally, such as in the center of histogram matching output for 2-HUH (first row) or the left side of cycleGAN output for 6-UTO (fifth row). A global error indicates a shift in the values of the output compared to the target and is attributed to the failure of the models to reconstruct the global contrast in the outputs accurately. On the other hand, local inaccuracies occur when the model can not capture the relative contrast between different areas of the brain or when it carries an undesired intensity gradient from the input to the output. In contrast, from the rightmost error column, we can observe that the overall difference to the target image is reduced noticeably in the output of the FP-cGAN model, and the error remains bounded throughout the whole brain compared to other methods. Additionally, by focusing on each column, we can observe that the heterogeneity of the output images of all three methods is lower than the input images. This is especially true for the output of the FP-cGAN model, suggesting better inter-dataset homogeneity of the harmonized images.

We present the numerical values of the calculated SSIM and PSNR similarity metrics in Table 2, which provides a comprehensive overview of the performance of each method. The table is divided into two horizontal sections, with each column representing the image harmonization results from a source site to a target site, except for the rightmost column (“AVERAGE”), which shows the averaged metrics over all source sites. The first row of each section corresponds to the pre-processed input images, while the remaining three rows represent the performance of the Hist-match, cycleGAN, and FP-cGAN methods. As can be seen, the FP-cGAN model achieves the highest average SSIM and PSNR values and improves the similarity to the target images from the input images in all five cases, demonstrating its effectiveness in MRI image harmonization. The table also highlights the comparative performance of each method: the SSIM metrics are highest in four out of five cases (shown in bold font), with cycleGAN yielding a slightly higher value by 0.1% in the 6-UTO site. The FP-cGAN model achieves the highest PSNR values in three cases. The Hist-match method improves both SSIM and PSNR in three sites. Furthermore, we observe that the PSNR metrics are improved by cycleGAN in four cases. In contrast, the SSIM metric is increased in only two cases when using cycleGAN. The rightmost column of Table 2 also reveals that the SSIM metrics between the input and target images are relatively high (93.14% on average) even before harmonization due to the fact that they belong to the same subject. However, the use of image harmonization methods leads to changes in these values: Hist-match increases the value to 93.59, while cycleGAN decreases it to 92.92. The FP-cGAN model exhibits the highest increase in average SSIM metrics to 94.15. This observation yields two key conclusions: first, an ineffective image harmonization method can have detrimental effects on the output image, even if theoretically relevant; second, as discussed in Section 3.5, it is not expected to witness a significant increase in similarity metrics when using effective MR image harmonization methods, due to the inherent closeness of input and target images. This is also because the target images are samples from a distribution rather than a concrete ground truth, which further limits the potential for improvement of similarity metrics.

**Table 2:** Average image-to-image similarity metrics SSIM and PSNR for different image harmonization methods. The metrics are calculated for the test subjects evaluated after transformation from different source sites, as indicated in the columns.

|  | 2-HUH | 3-HKH | 4-COI | 5-KUT | 6-UTO | AVERAGE |
| --- | --- | --- | --- | --- | --- | --- |
| SSIM: |  |  |  |  |  |  |
| Source (input) | 91.76 | 94.20 | 93.80 | 92.59 | 93.34 | 93.14 |
| Hist-match | 92.21 | 94.20 | 95.03 | 92.47 | 94.02 | 93.59 |
| cycleGAN | 91.14 | 94.05 | 94.35 | 91.01 | **94.06** | 92.92 |
| FP-cGAN (ours) | **93.58** | **94.51** | **95.23** | **93.48** | 93.96 | **94.15** |
| PSNR: |  |  |  |  |  |  |
| Source (input) | 27.00 | 29.63 | 25.72 | 27.02 | 26.06 | 27.09 |
| Hist-match | 26.93 | 29.45 | **32.01** | 27.18 | 28.49 | 28.81 |
| cycleGAN | 28.25 | 29.15 | 31.00 | 28.16 | **29.99** | 29.31 |
| FP-cGAN (ours) | **29.58** | **30.54** | 32.00 | **30.02** | 28.49 | **30.13** |

We also illustrate the calculated metrics for SSIM and PSNR in the form of boxplots in two subplots of Figure 5. Each group of columns belongs to the image harmonization results of the indicated source site. Within each group, from left to right, boxplots show the results for the similarity between the target image and the input image and outputs of Hist-match, cycleGAN, and FP-cGAN methods, illustrated in gray, blue, orange, and green colors, respectively, as indicated in the legend.

**Fig. 5:**
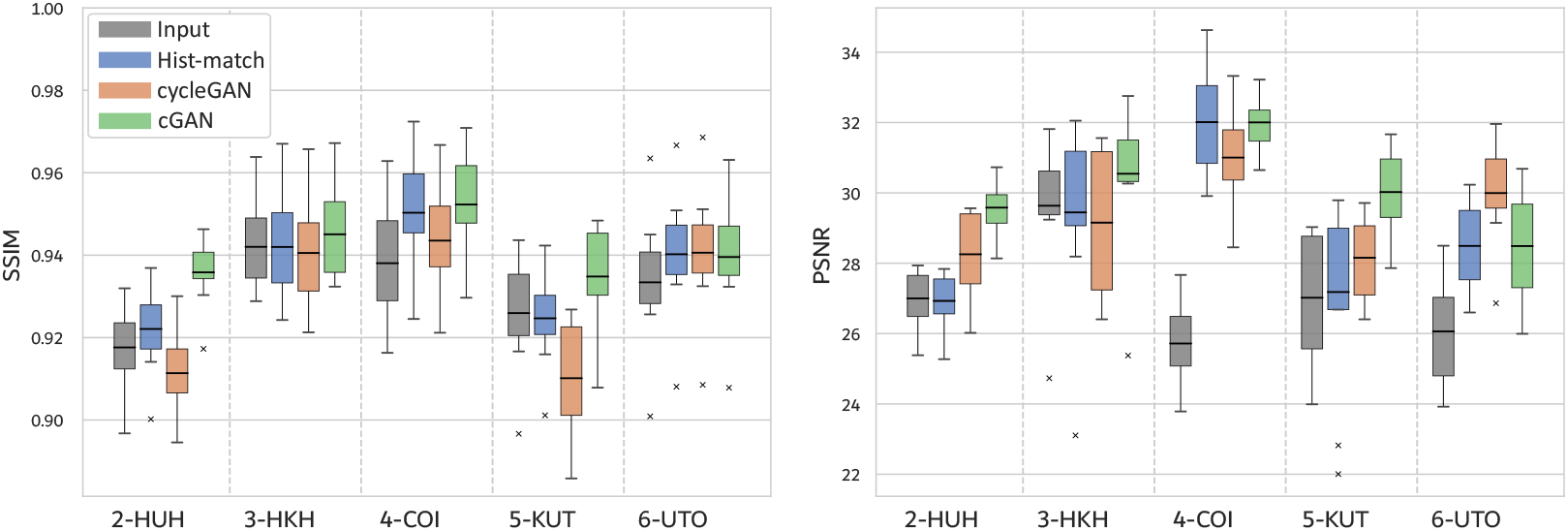
Boxplot results for the SSIM and PSNR metrics of histogram matching, cycle-GAN, and FP-cGAN are illustrated in blue, orange, and green, respectively. Each group of columns belongs to the harmonization results from the indicated site to the target site.

In addition to numerical analyses, we provide a few examples and visual interpretations in Figure 6 to demonstrate potential modes of inaccuracies in the outputs of cycleGAN and Hist-match methods. In this figure, from left to right, the first and second images are the input and target from the 6-UTO and 1-SWA sites. The third image is the output generated by the cycleGAN model with a rectangle that highlights a region of interest (ROI). The fourth and fifth images are the zoomed cycleGAN output and the error within the ROI. Similarly, the last two images are the output of the Hist-match method with an ROI and the output within that ROI. As can be observed in the fifth image (cycleGAN error ROI), the cycleGAN model has introduced minor rectangular artifacts in the output. These artifacts are barely discernible in the output image shown in the third and fourth images. They also do not have a significant impact on the calculated similarity metrics as Table 2 reports higher SSIM for the 6-UTO site compared to all other methods.

**Fig. 6:**
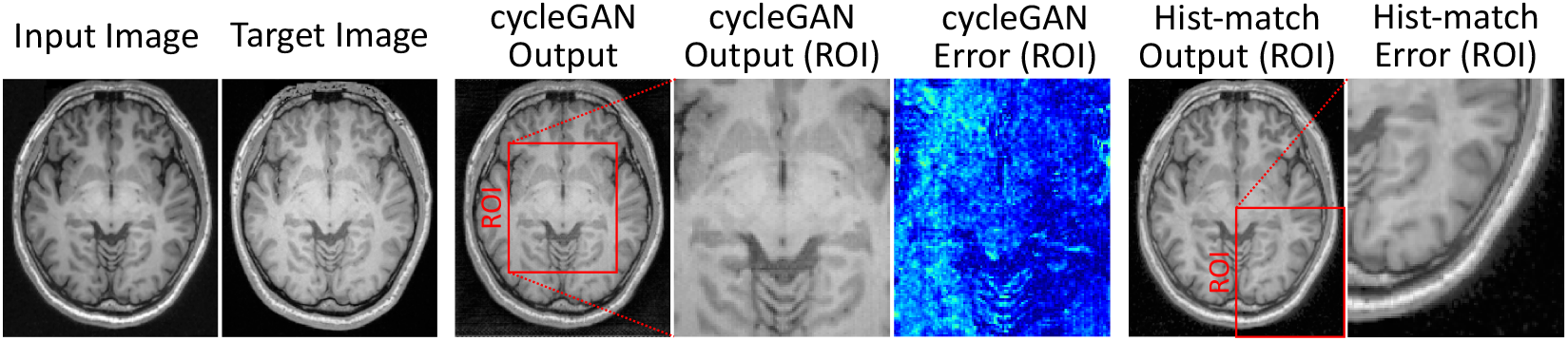
MR image harmonization examples from cycleGAn and Hist-match methods that demonstrate some common types of inaccuracies.

Figure 6 provides an example in which the target image is brighter than the input image. Naturally, the Hist-match method aims to increase the brightness of darker voxels to match the histogram of images. However, this can lead to an undesired increase in the noise seen in the rightmost image. This particular example can be mitigated by utilizing proper image processing methods. However, it demonstrates the potential risk of altering the voxel intensities of the tissues undesirably while matching the histograms. These challenges underscore the need for more robust and nuanced approaches, such as those introduced in the FP-cGAN model, to MRI image harmonization rather than merely relying on altering the histogram.

## 5 Conclusions and Future work

In this manuscript, we developed an FP-cGAN brain MR image harmonization model that transforms input images from various source sites to a specified target distribution. We utilized a generative adversarial scheme and constructed our model based on the distribution matching concept by minimizing the Wasserstein distance between the distribution of the transformed source images and images from the target site. We also introduced a regularization term to preserve the anatomical features of images during the harmonization process. We also showed that the proposed model is capable of providing an ensemble of transformed output images. This possibility is particularly desired in image harmonization problems, as any scanned image is considered a sample from a distribution. We also note that the proposed model does not require paired data for training, which is considered a critical property for an MRI image harmonization method.

We evaluated the performance of our FP-cGAN model and compared it against histogram matching and cycleGAN, two commonly performed methods in MRI image harmonization literature. We conducted our experiments using a dataset that contains traveling subjects for evaluation purposes, which enabled us to utilize image-to-image similarity metrics such as SSIM and PSNR. Based on our experiments, we concluded that the FP-cGAN model outperforms the other two methods and can effectively harmonize the input images to the target distribution.

Despite the above-mentioned advantages, this study has some limitations. For example, the model requires statistically representative source and target cohorts. However, in most neuroimaging studies, the goal is to harmonize MR images captured in a limited number of specific sites, and harmonizing MR images to a single reference image is not often necessary in practice. In future extensions of this work, we aim to investigate the effectiveness of the proposed model in harmonizing MR images used in a neuroimaging study. In this scenario, the evaluation process will involve investigating the statistical characteristics of distinct cohorts before and after harmonization

## Statements and Declarations

### Data Availability

No datasets were generated or analyzed during this study.

## Appendix

### A: Neural networks and hyper-parameters

We adopted the neural network architecture for the generator ***g*** and discriminator *d* discussed in [46] and applied necessary modifications. Notably, the discriminator in this work distinguishes fake versus real unpaired data. Therefore, the critic *d* does not require paired input data. We also removed the first level CIN block from the ResNet block. We set the hyperparameters *λ*_*gp*_ = 10 from the original work and *λ*_*gp*_ = 0.02 based on our experiments. We conducted our training experiments based on axial images with 240×200 resolution and a batch size of 16. During training, the generator weights were updated after every four updates of the critic weights. The model selection is based on an early stopping scheme using one subject from the traveling subject and SSIM similarity metrics. We used Adam optimizer [47] with amsgrad option [48] and set hyper-parameters *β*_1_ = 0.2, *β*_2_ = 0.7, the initial learning rate 3 × 10^*−*4^, and clipping norm of 10^6^.

## Footnotes

1 This dataset is part of the DecNef Project Brain Data Repository (https://bicr-resource.atr.jp/srpbsopen/), collected as part of the Japanese Strategic Research Program for the Promotion of Brain Science (SRPBS) supported by the Japanese Advanced Research and Development Programs for Medical Innovation (AMED).

